# WOX11-mediated adventitious lateral root formation modulates tolerance of Arabidopsis to cyst nematode infections

**DOI:** 10.1101/2023.08.11.553027

**Authors:** Jaap-Jan Willig, Nina Guarneri, Thomas van Loon, Sri Wahyuni, Ivan E. Astudillo-Estévez, Lin Xu, Viola Willemsen, Aska Goverse, Mark G. Sterken, José L. Lozano-Torres, Jaap Bakker, Geert Smant

**Affiliations:** Laboratory of Nematology, Wageningen University & Research, 6708 PB Wageningen, the Netherlands; National Key Laboratory of Plant Molecular Genetics, CAS Center for Excellence in Molecular Plant Sciences, Institute of Plant Physiology and Ecology, Chinese Academy of Sciences, Shanghai 200032, China; Cluster of Plant Developmental Biology, Cell and Developmental Biology, Wageningen University & Research, 6708 PB Wageningen, the Netherlands

**Author notes:** **Author for correspondence:** Geert Smant Postbus 8123, 6700ES, Wageningen. Instituto de Microbiología, Universidad San Francisco de Quito, Quito, Ecuador (Ivan E. Astudillo-Estévez). These authors have contributed equally (Jaap-Jan Willig, Nina Guarneri).

**Keywords:** Adventitious lateral root, COI1, damage, *de novo* root organogenesis, ERF109, *Heterodera schachtii*, LBD16, plasticity, root system architecture, WOX11, tolerance

## Abstract

The transcription factor *WUSCHEL-RELATED HOMEOBOX 11* (WOX11) in Arabidopsis initiates the formation of adventitious lateral roots upon mechanical injury in primary roots. Root-invading nematodes also induce *de novo* root organogenesis leading to excessive root branching, but it is not known if this symptom of disease involves mediation by WOX11 and if it benefits the plant. Here, we show with targeted transcriptional repression and reporter gene analyses in Arabidopsis that the beet cyst nematode *Heterodera schachtii* activates WOX11-adventitious lateral rooting from primary roots close to infection sites. The activation of WOX11 in nematode-infected roots occurs downstream of jasmonic acid-dependent damage signaling via *ETHYLENE RESPONSIVE FACTOR109*, linking adventitious lateral root formation to nematode damage to host tissues. By measuring different root system components, we found that WOX11-mediated formation of adventitious lateral roots compensates for nematode-induced inhibition of primary root growth. Our observations further demonstrate that WOX11-mediated rooting reduces the impact of nematode infections on aboveground plant development and growth. Altogether, we conclude that the transcriptional regulation by WOX11 modulates root system plasticity under biotic stress, which is one of the key mechanisms underlying tolerance of Arabidopsis to cyst nematode infections.

## Introduction

Soil-borne infections by cyst nematodes affect above- and below-ground plant development and growth, sometimes resulting in large yield losses in agriculture (Jones et al., 2013). Biotic stress induced by cyst nematodes in roots of host plants occurs at different stages of their infection cycle. Firstly, the infective second stage juveniles (J2s) invade host roots and migrate intracellularly through the epidermis and cortex, causing extensive damage to root tissue. Secondly, after becoming sedentary, cyst nematodes take up large amounts of plant assimilates during feeding from modified plant cells, which therefore develop strong metabolic sink activity (Gheysen and Mitchum, 2011; Jones et al., 2013; Bebber et al., 2014). As a response to nematode infections, plants remodel their root system by forming additional secondary roots (Goverse et al., 2000; Olmo et al., 2020; Willig et al., 2022; Guarneri et al., 2023). The *de novo* formation of secondary roots in response to endoparasitism by nematodes might be a mechanism to compensate for primary root growth inhibition caused by nematode infection (Guarneri et al., 2023). However, whether such a form of root system plasticity contributes to overall plant tolerance to cyst nematode infections remains to investigated.

Depending on where and how secondary roots are formed, they are either classified as lateral roots or adventitious lateral roots (Sheng et al., 2017). During post-embryonic development in Arabidopsis, periodic oscillations of auxin maxima at the root tip prime cells to form lateral roots that emerge in a regular acropetal pattern from the growing primary root (Fukaki and Tasaka, 2009; van den Berg et al., 2016). The emergence of lateral roots is controlled by AUXIN RESPONSE FACTOR (ARF)7 and ARF19, which directly regulate *LATERAL ORGAN BOUNDARIES DOMAIN* (*LBD)16* and other *LBD* genes (Okushima et al., 2007). In contrast, adventitious lateral roots do not follow an acropetal pattern as they emerge in between and opposite of existing lateral roots. Moreover, adventitious lateral roots emerge in response to tissue damage, and their formation is regulated by a separate pathway mediated by the transcription factor WUSCHEL-RELATED HOMEOBOX (WOX)11 (Liu et al., 2014; Hu and Xu, 2016; Sheng et al., 2017). After cutting the primary root, local accumulation of auxin activates *WOX11* transcriptional activity through auxin response elements in its promotor region (Liu et al., 2014). Subsequently, WOX11 induces the expression of *LBD16* but also the expression of other *WOX* genes (Hu and Xu, 2016; Sheng et al., 2017). Ultimately, this leads to the *de novo* formation of secondary roots close to the injury site (Cai et al., 2014; Liu et al., 2014; Hu and Xu, 2016; Sheng et al., 2017). Cyst nematode infection in primary roots of Arabidopsis triggers the formation of secondary roots which does not follow an acropetal patterning (Guarneri et al., 2022). Instead, secondary roots often form clusters at nematode infection sites. As to whether the formation of these secondary roots depends on the WOX11-mediated pathway and whether they should thus be classified as adventitious lateral roots is still a knowledge gap.

We have recently demonstrated that the formation of secondary roots near nematode infection sites involves damage-induced jasmonic acid (JA) signalling (Guarneri et al., 2023). Tissue damage caused by intracellular migration of infective juveniles of *H. schachtii* induces the biosynthesis of JA, which activates the transcription factor *ETHYLENE RESPONSIVE FACTOR* (ERF)109 via the JA receptor *CORONATINE INSENSITIVE* (COI)1. ERF109, in turn, can trigger local biosynthesis of auxin by directly binding to the promoters of auxin biosynthesis genes *ASA1* and *YUC2* (Cai et al., 2014). Indeed, our data showed that COI1/ERF109-mediated formation of secondary roots from nematode-infected primary roots depends on local biosynthesis and accumulation of auxin (Guarneri et al., 2023). WOX11-mediated formation of adventitious lateral roots upon root injury also involves local accumulation of auxin (Liu et al., 2014). However, it remains to be demonstrated if WOX11 becomes activated by COI1- and ERF109-mediated damage signalling in nematode-infected roots.

Several recent reports in the literature point at a role for WOX11-mediated root plasticity in modulating plant responses to abiotic stresses. For instance, *WOX11*, designated as *PagWOX11/ WOX12a*, in poplar mediates changes in root system architecture in response to drought and salt stress (Wang et al., 2020; Wang et al., 2021). Overexpression and dominant repression of this gene in poplar plants alters the number of adventitious roots formed under high saline conditions (Liu et al., 2022). Likewise, the loss-of-function mutant *wox11* in rice exhibits reduced root system development in response to drought as compared to wildtype plants (Cheng et al., 2016). Based on these findings, WOX11-mediated root plasticity is thought to enhance plant tolerance to abiotic stress. However, whether WOX11-mediated root plasticity is also involved in mitigating the impact of biotic stresses on the root system is not known.

In this study, we first addressed whether cyst nematode-induced secondary roots qualify as damage-induced adventitious lateral roots. Hereto, we monitored *de novo* secondary root formation in Arabidopsis seedlings of the double mutant *arf7/arf19* and the WOX11 transcriptional repressor mutant *35S:WOX11-SRDX* in the *arf7/arf19* background (Hiratsu et al., 2003) infected with *H. schachtii.* Next, we asked whether the regulation of *WOX11* in nematode-infected Arabidopsis roots occurs downstream of JA-dependent damage signalling through COI1 and ERF109. To answer this question we performed a time course experiment measuring *pWOX11::GFP* expression with confocal microscopy in wild-type, *coi1-2*, and *erf109* infected mutant seedlings. Third, we assessed if WOX-11-mediated root system plasticity compensates for the inhibition of primary root growth upon cyst nematode infection. For this, we measured different components of root system architecture of nematode-infected WOX11 transcriptional repressor mutant and wildtype Arabidopsis plants. Last, we tested if WOX11-mediated root system plasticity contributes to the overall tolerance of Arabidopsis to cyst nematode infections. To this end, we compared the aboveground plant growth and development of cyst nematode-infected *35S:WOX11-SRDX* mutants and wildtype Arabidopsis for a period of three weeks after inoculation. Based on our data, we propose a model wherein the formation of WOX11-mediated adventitious lateral roots enhances tolerance of Arabidopsis to biotic stress by cyst nematode infections.

## Materials and Method

### Plant material and culturing

The Arabidopsis (Arabidopsis thaliana) lines wild-type Col-0 (N60.000), *35S:WOX11-SRDX/arf7-1/19-1*, *arf7-1/19-1*, *LBD16pro:LBD16-GUS* and *35S:WOX11-SRDX/LBD16pro:LBD16-GUS* (Sheng et al., 2017), *pWOX11::GFP*, *pWOX11::GFP-coi1-2*, *pWOX11::GFP-erf109*, *coi1-2* and *erf109* were used. For in vitro experiments, seeds were vapor sterilized for 3-4 hours using a mixture of hydrochloric acid (25%) and sodium hypochlorite (50 g/L). Finally, sterile seeds were stratified for 4 days at 4 °C, after which they were sown on square Petri dishes (120x120 mm) containing modified Knop medium (Sijmons et al., 1991) in a growth chamber with a 16-h-light/8-h-dark photoperiod at 21°C. For in vivo pot experiments, seeds were stratified for 4 days and sown on silver sand in 200 mL pots. Seedlings were grown at 19 °C and 16-h-light/8-h-dark conditions with LED light (150 lumen), as previously described in (Willig et al., 2023).

### Hatching and sterilization of *Heterodera schachtii*

*H. schachtii* cysts (Woensdrecht population from IRS, the Netherlands) were separated from sand of infected *Brassica oleracea* plants as previously described (Baum et al., 2000). Cysts were transferred into a clean Erlenmeyer containing water with 0.02% sodium azide. This mixture was gently stirred for 20 min. Later, sodium azide was removed by washing with tap water. Cysts were then incubated for 4-7 days in a solution containing 1.5 mg/mL gentamycin sulfate, 0.05 mg/mL nystatin and 3 mM ZnCl_2_. Hatched J2s were purified by centrifugation on a 35% sucrose gradient, transferred to a 2 mL Eppendorf tube and surface sterilized for 15 minutes in a solution containing 0.16 mM HgCl_2_, 0.49 mM NaN_3_, and 0.002% (v/v) Triton X-100. After washing the J2s three times with sterile tap water, *H. schachtii* J2s were re-suspended in a sterile 0.7% Gelrite (Duchefa Biochemie, Haarlem, the Netherland) solution. A similar concentration of Gelrite solution was used as mock treatment.

For *in vivo* pot experiments, J2s were hatched and collected in a similar way as described above. Non-sterile J2s were purified by centrifugation on a 35% sucrose gradient and washed three times with tap water. Nematodes were resuspended in tap water for specific inoculation densities.

### Quantifying root system architecture of nematode-infected Arabidopsis

Seven-day-old *35S:WOX11-SRDX/arf7-1/19-1* and *arf7-1/19-1* Arabidopsis seedlings were inoculated with either 90 *H. schachtii* J2s or a mock solution. Root architecture was inspected at 7 dpi using an Olympus SZX10 binocular with a 1.5x objective and 2.5x magnification. Scans were made of whole seedlings using an Epson Perfection V800 photo scanner. Pictures of nematode infections were taken with a AxioCam MRc5 camera (Zeiss) and the ZEN 3.2 blue edition software (Zeiss).

Nine-day-old *35S:WOX11-SRDX* and wild-type Col-0 seedlings, grown on 120x120 mm square Petri dishes were inoculated with 0 (mock), 0.5, 1.0, 2.5, 5.0, and 7.5 *H. schachtii* J2s per mL of modified Knop medium as previously described (Guarneri et al., 2023). Inoculations were done with two 5 µl drops that were pipetted at opposite sides of each seedling while keeping the petri dishes vertical. At 7 dpi, scans were made of whole seedlings using an Epson Perfection V800 photo scanner. The architecture (i.e., total root length, primary root length, total secondary root length) was measured using the WinRHIZO package for Arabidopsis (WinRHIZO pro2015, Regent Instrument Inc., Quebec, Canada). The number of root tips was counted manually based on the scans.

### Acid fuchsine staining of nematodes

Nematodes within the roots were stained with acid fuchsin and counted as previously described (Warmerdam et al., 2018). For comparisons between genotypes, the background effect of the mutation on the root architecture was corrected by normalizing each measured root architecture component in infected seedlings to the median respective component in mock-inoculated roots.

### Histology and brightfield microscopy

Four-day-old Arabidopsis seedlings were inoculated with 20 *H. schachtii* J2s or a mock solution. For histochemical staining of β-glucuronidase (GUS) activity, seedlings were incubated in a GUS staining solution (1 mg/mL X-GlcA in 100 mM phosphate buffer pH 7.2, 2 mM potassium ferricyanide, 2 mM potassium ferrocyanide, and 0.2 % Triton X-100) at 37 °C (Zhou et al., 2019) for 3 hours. Stained seedlings were mounted in a chloral hydrate clearing solution (12 M chloral hydrate, 25% glycerol) and inspected with an Axio Imager nM2 light microscope (Zeiss) via a 20x objective. Differential interference contrast (DIC) images were taken with an AxioCam MRc5 camera (Zeiss) and the ZEN 3.2 blue edition software (Zeiss).

### Confocal laser microscopy of single *H. schachtii* infection sites

Four-day-old Arabidopsis seedlings were inoculated with roughly five sterile *H. schachtii* J2s in 10 µL 0.7% Gelrite. Single nematode infection sites were selected for observation at 2, 3, 4, and 7 dpi. Infection sites were inspected using a Zeiss LSM 710 confocal laser scanning microscope and a 40x objective. After a single infection site was located, a Z-stack of ten 13 μm-slices was made. Z-stacks were taken using the ZEN 2009 software (Zeiss). The imaging settings in ZEN 2009 were as follows: Laser 488 at 50%, Pinhole 41.4 µm, eGFP 645 nm, TPMT 217 nm. Z-stacks were processed with ImageJ Version 1.53 to quantify the fluorescence integrated density.

The post-processing in ImageJ of one individual image was as follows: Firstly, an auto-scaled compressed-hyper-Z-stack was created of the 10 layers made with the confocal microscope by using the Z-compression function at max intensity (Supplemental Figure S4). Secondly, a duplicate of the original Z-stack was created, and a Gaussian filter with a sigma value of 2.0 was applied to this duplicate. This duplicate was subtracted from the original image by using the image calculator function. Thirdly, the image threshold limits were set to a specific range ranging from 0 to 100 depending on the quality of the image. The same threshold limits were applied on all images that were taken on the same day. Lastly, the particles were analysed using Analyse Particles at size 0-Infinity and circularity 0.00-1.00.

### High throughput analysis of the green canopy area of nematode-infected Arabidopsis plants

Plants were imaged and analyzed as previously described (Willig et al., 2023). Prior to sowing, pots, were filled with silver sand, covered with black coversheets, and were watered with Hyponex (1.7 mM/L NH_4_^+^, 4.1 mM/L K^+^, 2 mM/L Ca_2_^+^, 1.2 mM/L Mg_2_^+^, 4.3 mM/L NO_3_^-^, 3.3 mM/L SO_4_^2-^, 1.3 mM/L H2PO_4_^-^, 3.4 µm/L Mn, 4.7 µm/L Zn, B 14 µm/L, 6.9 µm/L Cu, 0.5 µm/L Mo, 21 µm/L Fe, pH 5.8) for five minutes. Seven days after sowing, seedlings were watered again for five minutes. Nine-day-old seedlings were inoculated with increasing densities of *H. schachtii* (0 to 10 juveniles per g dry sand). For our experiments we did not use a blocking design as it would greatly increase the chance for error when manually inoculating plants. Every hour, pictures were taken of the plants (15 pictures per day) for a period of 21 days. At the end of the experiment, colour corrections were done using Adobe Photoshop (Version: 22.5.6 20220204.r.749 810e0a0 x64). The surface area of the rosette was determined using a custom-written ImageJ macro (ImageJ 1.51f; Java 1.8.0_321 [32-bit]) and Java was used to make GIFs.

### Plant growth analysis and tolerance modelling using a high-throughput phenotyping platform

To analyse the growth data of the plants obtained from the high-throughput platform, we followed the same approach and used the same functions as in our previously published analytical pipeline (Willig et al., 2023); available via Gitlab: https://git.wur.nl/published_papers/willig_2023_camera-setup).

In short, the measurement used was the median daily leaf area (cm^2^), calculated from the 15 daily measurements. We used log_2_-transformed data, where the rate of growth was determined per day per plant by (equation 1)

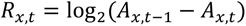

where *R_x,t_* is the transformed growth rate of plant *x* at day *t* from day *t-1* to day *t* based on the median Green canopy area *Ax,t*.

The tolerance limit was modelled using a previously described method based on fitting growth models (Willig et al., 2023). Here we fitted a logistic growth model using the *growthrates* package on the median daily leaf area A_t_ (cm^2^) (equation 2),

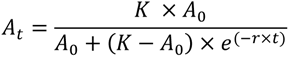

where *K* is the maximum green canopy area (cm^2^), *A_0_* is the initial canopy area (cm^2^), and *r* is the intrinsic growth rate (d^-1^), which were determined as a function of time *t* (d) (p < 0.1).

Based on the relation between *K* and density we could identify the tolerance limit (equation 3)

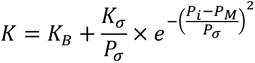

where *P_i_* is the initial nematode density in nematodes per gram soil, *K_B_* is the basal canopy size, *K_σ_* is the normalized maximum canopy area that can be achieved over the *P_i_* range, *P_σ_* is the deviation around the nematode density allowing maximum growth, *P_M_* is the nematode density at which maximum growth is achieved. We modelled the parameter values using *nls* and extracted confidence intervals using the *nlstools* package (Baty et al., 2015). The tolerance limit, 2\**P_M_,* could such be determined (as in (Willig et al., 2023)).

### Statistical analyses

Statistical analyses were performed using the R software version 3.6.3 (Windows, x64). The R packages used are *tidyverse* (https://CRAN.R-project.org/package=tidyverse), *ARTool* (https://CRAN.R-project.org/package=ARTool) and *multcompView* (https://CRAN.R-project.org/package=multcompView).

Correlation between variables was calculated using Spearman Rank-Order Correlation coefficient. For binary data, significance of the differences between proportions was calculated by a Pairwise Z-test. For normally distributed data, significance of the differences among means was calculated by ANOVA followed by Tukey’s HSD test for multiple comparisons. A non-parametric pairwise Wilcoxon test followed by false discovery rate correction for multiple comparisons was used for data with other distributions and one grouping factor. For the high-throughput platform data we used the Wilcoxon test as implemented in the *ggpubr* package (https://cran.r-project.org/web/packages/ggpubr/index.html). The confidence interval of the inoculum density-response curves was calculated by LOESS regression (as per default in *geom_smooth*) in R.

## Results

### Cyst nematodes induce the formation of adventitious lateral roots

Our earlier work showed that *H. schachtii* induces the *de novo* formation of secondary roots between or across fully developed lateral roots near nematode infection sites (Guarneri et al., 2023). Here, we hypothesized that these secondary roots are adventitious lateral roots, the formation of which depends on WOX11-mediated transcriptional regulation (Fig. 1A). To test this hypothesis, we inoculated *H. schachtii* on the lateral root-deficient *arf7-1/19-1* double mutant, which is unable to form acropetal lateral roots, and the transcription repressor mutant *35S:WOX11-SRDX/arf7-1/19-1* (Hiratsu et al., 2003), which is unable to form neither acropetal nor adventitious lateral roots (Fig. 1B-D). Importantly, we observed no difference in the number of nematodes per plant between *arf7-1/19-1* and *35S:WOX11-SRDX/arf7-1/19-1* (Fig. 1C), indicating that both Arabidopsis lines were exposed to similar levels of biotic stress. However, while *H. schachtii* induced the formation of secondary roots on *arf7-1/19-1* mutant line (Fig. 1B, D and Supplemental Fig. S1), no secondary roots emerged from nematode-infected roots of the *35S:WOX11-SRDX/arf7-1/19- 1* mutant. From this, we concluded that the induction of secondary roots by *H. schachtii* is mediated by WOX11 and that these secondary roots therefore qualify as adventitious lateral roots.

**Figure 1:**
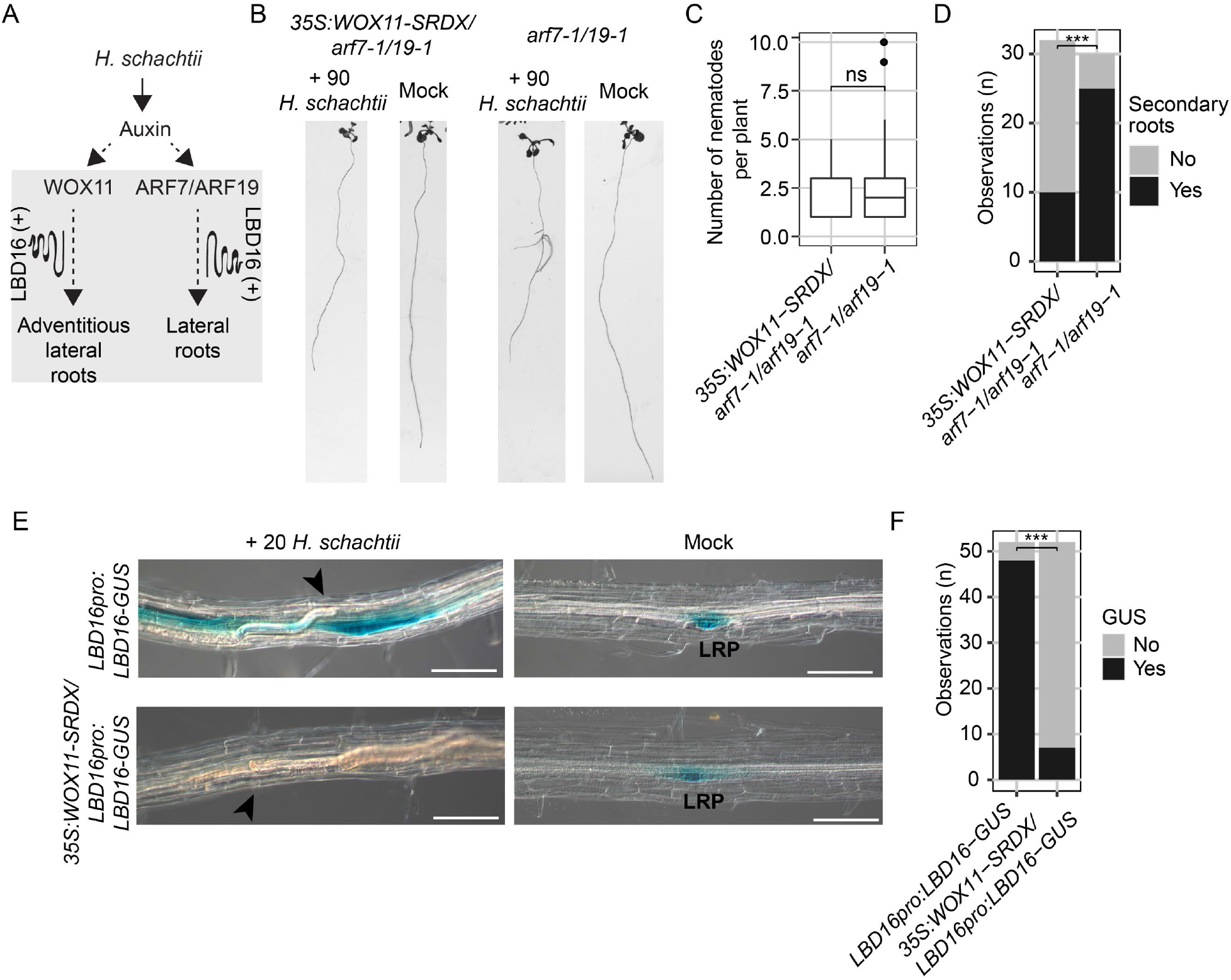
*Heterodera schachtii* induces adventitious lateral root formation in a WOX11- and LBD16-dependent manner. **A)** Schematic diagram of *H. schachtii*-and WOX11-mediated adventitious lateral root emergence. Grey area indicates the tested part of the pathway. Curling line and ‘+’ indicate involvement of multiple proteins, including LBD16. **B-D)** Seven-day old *35S:WOX11- SRDX/arf7-1/19-1* and *arf7-1/arf19-1* mutant seedlings were inoculated with 90 *H. schachtii* juveniles or mock. At 7dpi, scans were made of the root system. **B)** Representative pictures of *35S:WOX11-SRDX/arf7-1/19-1* and *arf7-1/arf19-1* mutant seedlings inoculated with 90 *H. schachtii* or with mock solution. **C)** Number of juveniles that invaded the primary roots. **D)** Number of seedlings that show secondary roots (Yes) that are associated with *H. schachtii* infection sites or no secondary roots at all (No). Data from three independent biological repeats of the experiment was combined. Statistical significance was calculated by a Pairwise Z-test n=30-32, ***: p<0.001). **E-F)** Four-day-old Arabidopsis seedlings expressing the *LBD16pro:LBD16-GUS* and *35S:WOX11- SRDX/LBD16pro:LBD16-GUS* reporters were inoculated with 20 *H. schachtii* juveniles. At 4dpi, GUS expression was stained for 3 hours and seedlings were imaged **e)** *LBD16pro:LBD16-GUS* and *35S:WOX11-SRDX/LBD16pro:LBD16-GUS* expression at nematode infection sites in roots. Black arrowheads indicate the nematode head. LRP indicates lateral root primordia. Scale bar = 100 µm. **F)** Number of observations with (Yes) or without (No) GUS staining at the nematode infection site in roots of wild-type Col-0 seedlings. Data from three independent biological repeats of the experiment were combined. Statistical significance was calculated by a Pairwise Z-test n=52, ***: p<0.001).

WOX11-mediated formation of adventitious lateral roots from primary roots of Arabidopsis involves the downstream transcriptional activation of *LBD16* (Fig. 1A) (Sheng et al., 2017). To test if WOX11 activates *LBD16* in nematode-infected Arabidopsis roots, we monitored the expression of *LBD16* fused to *GUS* in wildtype (*LBD16pro:LBD16-GUS*) and *35S:WOX11-SRDX*(*35S:WOX11-SRDX/LBD16pro:LBD16-GUS*) seedlings inoculated with *H. schachtii* (Fig. 1E and F). We found that *LBD16* was highly expressed in nematode infection sites in the wildtype, but not in the *35S:WOX11-SRDX* background. This demonstrates that *H. schachtii* activates *LBD16* expression in a WOX11-dependent manner. Based on these observations, we concluded that cyst nematode infections activate the WOX11/LBD16-mediated pathway to form adventitious lateral roots from primary roots.

### Emergence of adventitious lateral roots correlates with damage in primary roots

Previously, we showed that increasing nematode inoculation densities result in more tissue damage in Arabidopsis leading to a higher number of secondary roots emerging from infected primary roots (Guarneri et al., 2023). To test the hypothesis that WOX11 mediates this quantitative relationship between inoculation density and the number of secondary roots emerging from cyst nematode-infected primary roots (Fig. 2A), we inoculated nine-day-old seedlings of *35S:WOX11-SRDX* and wildtype plants with increasing densities of *H. schachtii* (Fig. 2B). At 7 dpi, the number of nematodes that had successfully penetrated the roots was counted after staining with acid fuchsin (Fig. 2C). The number of infective juveniles in *35S:WOX11-SRDX* plants by inoculation density was significantly higher compared to wild-type Col-0 plants. This indicates that the transcriptional regulation by WOX11 in wild-type Arabidopsis plants reduces susceptibility to penetration by *H. schachtii*. Next, we counted the number of secondary roots to determine whether this correlates with the number of nematodes inside the roots. It should be noted that uninfected *35S:WOX11-SRDX* plants have more secondary roots than wild-type Col-0 plants (Supplemental Fig. S2). To correct for this background effect of the SRDX-transcriptional repressor construct on root system architecture, we normalized the total number of secondary roots in infected seedlings to the average respective number in uninfected seedlings (Fig. 2D). As expected, after normalization, the number of secondary roots emerging from primary roots increased with the number of successful invasions of *H. schachtii* in wild-type Arabidopsis. However, no such correlation was observed in *35S:WOX11-SRDX* plants. We therefore concluded that the density-dependent adaptations in root system architecture to increasing levels of damage in nematode-infected roots are brought about by WOX11-mediated formation of adventitious lateral roots.

**Figure 2:**
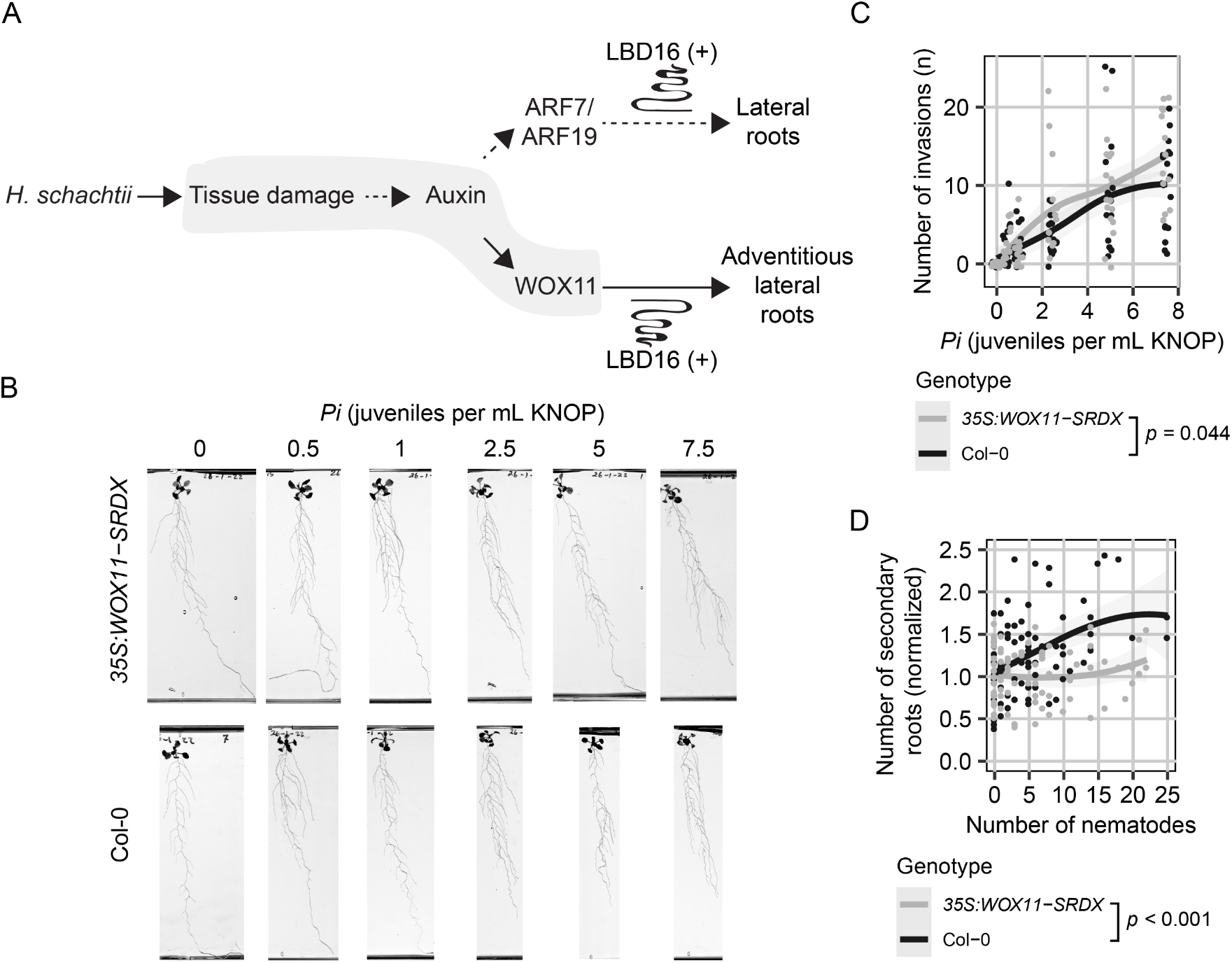
WOX11 is required in *Heterodera schachtii* induced adventitious lateral root formation in a density-dependent manner. **A)** Schematic diagram of *H. schachtii*-and WOX11-mediated adventitious lateral roots emergence. Grey area indicates the tested part of the pathway. Curling line and ‘+’ indicate the involvement of multiple proteins, including LBD16. **B-D)** Nine-day-old *35S:WOX11-SRDX* and wild-type Col-0 seedlings were inoculated with nematode densities (*Pi*) ranging from 0-7.5 *H. schachtii* J2s (mL modified KNOP media). Roots were scanned and nematodes were counted after acid fuchsin staining at 7 dpi. **B)** Representative images of Arabidopsis root systems at 7dpi. **C)** Number of nematodes that successfully penetrated the roots per plant. **D)** Secondary roots formed per number of nematodes inside the roots. The total number of secondary roots of infected seedlings was normalized to the median respective component in mock-inoculated roots. Data from two independent biological repeats of the experiment were combined. Significance of differences between genotypes was calculated by analysis of variance (n=14-18). Grey area indicates the 95% confidence interval.

### COI1 and ERF109 modulate damage-induced activation of *WOX11* at nematode infection sites

*De novo* formation of secondary roots on nematode-infected primary roots of Arabidopsis is mediated by damage-induced activation of JA signaling via COI1 and ERF109 (Guarneri et al., 2023). In this study we tested COI1 and ERF109 are required for the regulation of *WOX11* in nematode-infection sites (Fig. 3A). Hereto, we imaged nucleus-localized *pWOX11::GFP* expression within single-nematode infection sites in the *coi1-2* and *erf109* mutants and wild-type Col-0 at 2, 3, 4, and 7 dpi (Fig. 3 and Supplemental Fig. S3). Cyst nematode infection typically causes tissue autofluorescence in Arabidopsis roots (Hoth et al., 2005). To filter out this autofluorescence from the fluorescent signal emitted by the GFP construct, we subtracted a Gaussian blurred image from the original images (Supplemental Fig. S4). Hereafter, we observed a gradual increase in the *pWOX11::GFP*-derived fluorescent signal in nematode infection sites over time in *coi1-2, erf109,* and wild-type Col-0 (Fig. S3A-F), with wild-type Col-0 showing the strongest increase (Supplemental Fig. S3F). For instance, at 4 dpi, wild-type Col-0 plants showed significantly more nuclear GFP fluorescence in and around nematode feeding sites compared to *coi1-2* and *erf109* in the processed images (Fig. 3B-D). We, therefore, concluded that two key components of the damage-induced JA signalling pathway, COI1 and ERF109, modulate *WOX11* expression in infection sites of *H. schachtii* in Arabidopsis.

**Figure 3:**
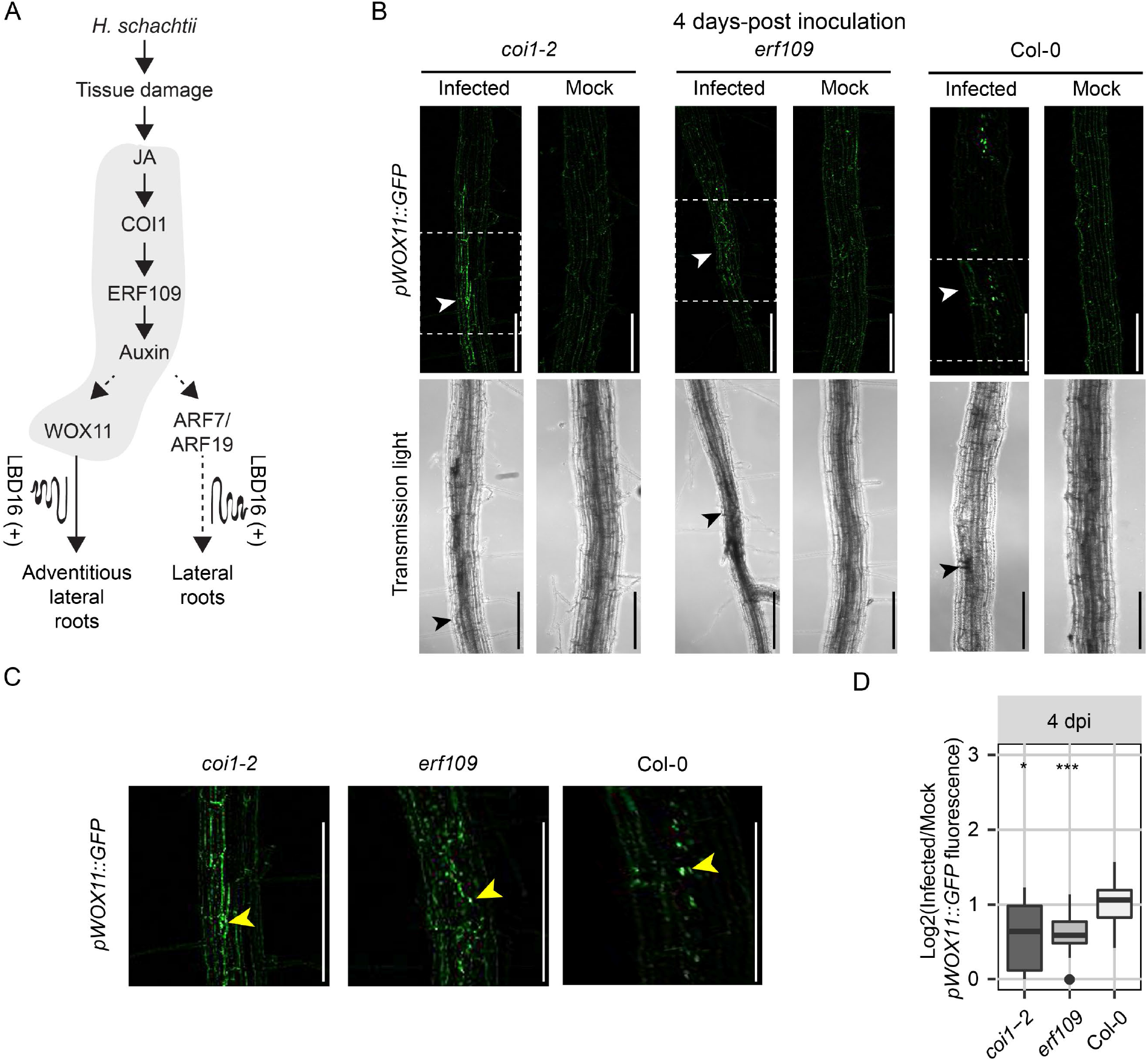
COI1 and ERF109 modulate *WOX11* expression upon *H. schachtii* infection. **A)** Schematic diagram of *H. schachtii*-and WOX11-mediated adventitious lateral root emergence. Grey area indicates the tested part of the pathway. Curling line and ‘+’ indicate involvement of multiple proteins, including LBD16. **B-C)** Four-day-old Arabidopsis seedlings were either inoculated with 10 *H.* schachtii second-stage juveniles (J2s) or mock-inoculated. At 4 dpi, seedlings were mounted in water and then imaged using a fluorescent confocal microscope. Single-nematode infection sites were selected for observation. Images are original. **B)** Representative pictures of infected and mock-inoculated seedlings expressing the *pWOX11::GFP* construct with nuclear localization signal in either wild-type Col-0, mutant *coi1-2*, or mutant *erf109* background at 4 dpi. To make the fluorescence more visible, the brightness was enhanced for all the representative pictures in the same way. **C)** Zoomed parts of original images fluorescent signal that are indicated by dashed white box in panel **(B)**. Yellow arrowhead indicates true fluorescent signal of *pWOX11::GFP* in the nucleus. **D)** Quantification of *pWOX11::GFP* fluorescent intensity induced by infection in wild-type Col-0, *coi1-2*, and *erf109* roots. Values represent log2 of the fluorescence ratio between the GFP integrated density of infected and noninfected roots. Scale bar: 200 µm. Data from three independent biological repeats of the experiment were combined. Significance of differences between fluorescent intensities in Co-0, *coi1-2*, and *erf109* per timepoint was calculated by a Wilcoxon Rank Sum test. ns = not significant, *p< 0.05, **p< 0.01, ***p<0.001 (n=15).

### Formation of adventitious lateral roots compensates for nematode-induced primary root growth inhibition

Next, we asked whether WOX11-mediated adventitious lateral roots formation compensates for the inhibition of primary root growth due to nematode infections (Fig. 4A). To this end, we quantified root system architecture components (i.e., total root length, primary root length, total secondary root length, and average secondary root length) of nematode-infected roots of both *35S:WOX11-SRDX* and wildtype Col-0 plants (Fig. 2). Initially, we noticed that our measurements of root system architecture components followed a parabolic function with the minimum values at the infection rate of 15 juveniles per root, suggesting the existence of two density dependent counteracting mechanisms (Supplemental Fig. S5). We, therefore, analysed our data for the lower (Fig. 4) and higher infection rates separately (Supplemental Fig. S5). For plants infected with 0 to 15 juveniles per root, we found that the total root length was significantly more reduced by nematode infection in *35S:WOX11-SRDX* mutant plants than in wild-type Col-0 plants (Fig. 4B and D). Interestingly, the growth of the primary root was not different between *35S:WOX11-SRDX* mutant plants and wild-type plants upon infection with cyst nematodes (Fig. 4C). However, the total length of the secondary roots of nematode-infected *35S:WOX11-SRDX* mutant plants was significantly smaller as compared to wild-type Col-0 plants (Fig. 4D). As the average secondary root length did not significantly differ between *35S:WOX11-SRDX* and wild-type Arabidopsis plants, WOX11 affects the root system architecture by increasing the number of secondary roots but not by extending secondary root growth (Supplemental Fig. 5E). For plants infected with 15 to 25 juveniles per plant, we observed no significant differences for the total root length (Supplemental Fig. S5B) between wild-type Col-0 and *35S:WOX11- SRDX*. Likewise, we found no differences in the primary root length (Supplemental Fig. S5C), total secondary root length (Supplemental Fig. S5D), and average secondary root length (Fig. S5E). Based on our analyses, we concluded that WOX11-mediated formation of adventitious lateral roots compensates for nematode-induced inhibition of primary root growth at lower infection rates.

**Figure 4:**
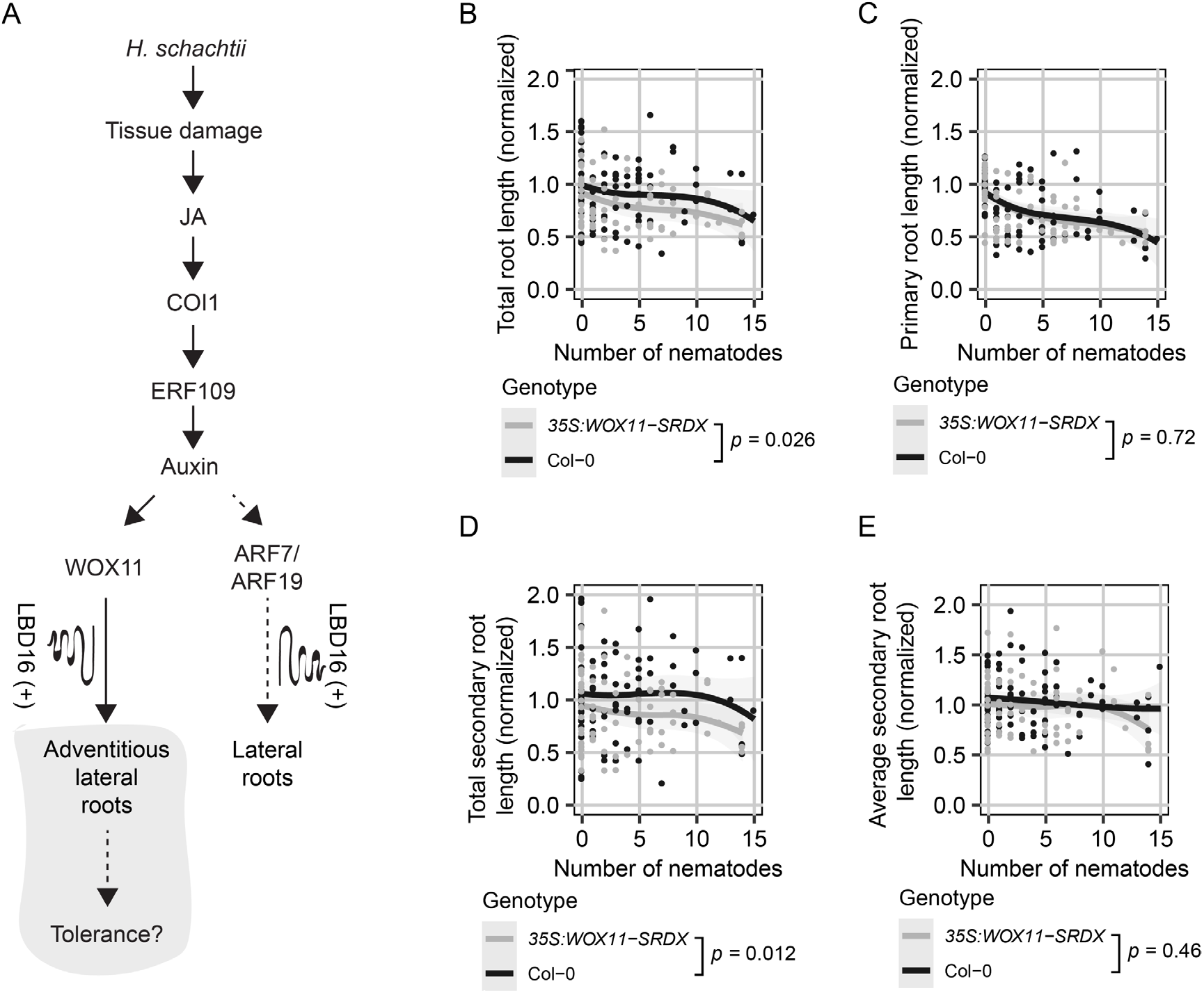
Formation of adventitious lateral roots compensates for nematode-induced primary root growth inhibition. **A)** Schematic diagram of *H. schachtii*-and WOX11-mediated adventitious lateral root emergence. Grey area indicates the tested part of the pathway. Curling line and ‘+’ indicate involvement of multiple proteins, including LBD16. **B-D)** Nine-day old *35S:WOX11-SRDX* and wild-type Col-0 seedlings were inoculated densities (*Pi*) ranging from 0-7.5 *H. schachtii* J2s (mL modified KNOP media). Roots were scanned and nematodes were counted after fuchsine staining at 7 dpi. Root architectural components of infected seedlings were normalized to the median respective component in mock-treated roots. Data of two independent biological repeats of the experiment was combined. **B)** Representative images of Arabidopsis root system at 7dpi. **B)** Total root length per number of nematodes inside the roots. **C)** Primary root length per number of nematodes inside the roots. **D)** Total secondary root length per number of nematodes inside the roots. **E)** Average secondary root length per number of nematodes inside the roots. Data from two independent biological repeats of the experiment were combined. Significance of differences between genotypes was calculated by analysis of variance (n=14-18). Grey area indicates the 95% confidence interval of the LOESS fit.

### WOX11 modulates tolerance to cyst nematode infections

The growth of the green canopy area over time reflects the tolerance of Arabidopsis to biotic stress by root-feeding cyst nematodes (Willig et al., 2023). To assess if WOX11-mediated *de novo* formation of adventitious lateral roots modulates tolerance of Arabidopsis to cyst nematode infection, we monitored the growth of the green canopy area of *35S:WOX11-SRDX* mutant and wild-type Col-0 seedlings for a period of 21 days after inoculation with different numbers of *H. schachtii* (Fig. 5A and B). At the end of the experiment, the green canopy area of the *35S:WOX11-SRDX* mutant was smaller at higher inoculation densities of *H. schachtii* as compared to wild-type Col-0 plants (Fig. 5C and D). Notably, the first significant reduction in green canopy area of *35S:WOX11-SRDX* plants by nematode infection was observed at inoculation densities between *P_i_* 2.5 and 5 J2s per gram sand, while in wild-type Col-0 plants we observed a first significant reduction in green canopy area at *P_i_* 7.5 J2s per gram sand. To quantify more exactly the difference in tolerance of *35S:WOX11-SRDX* and wildtype Col-0 plants, we fitted the growth rates of individual plants (Supplemental Fig. S6 and S7) to a logistic growth model. From this, we calculated the maximum projected green canopy area and determined the tolerance limit with 95% confidence interval (95% CI) (Fig. 5E). The relationship between maximum canopy area *K* and the *P_i_* fitted a Gaussian curve, based on which we estimated the tolerance limit for *35S:WOX11-SRDX* at *P_i_* = 2.25 (95% CI: 0.67-3.83) and for wild-type Col-0 at *P_i_* = 4.84 (95% CI: 3.8-5.89). This difference in tolerance limits led us to conclude that WOX11 modulates tolerance of Arabidopsis to cyst nematode infections.

**Figure 5:**
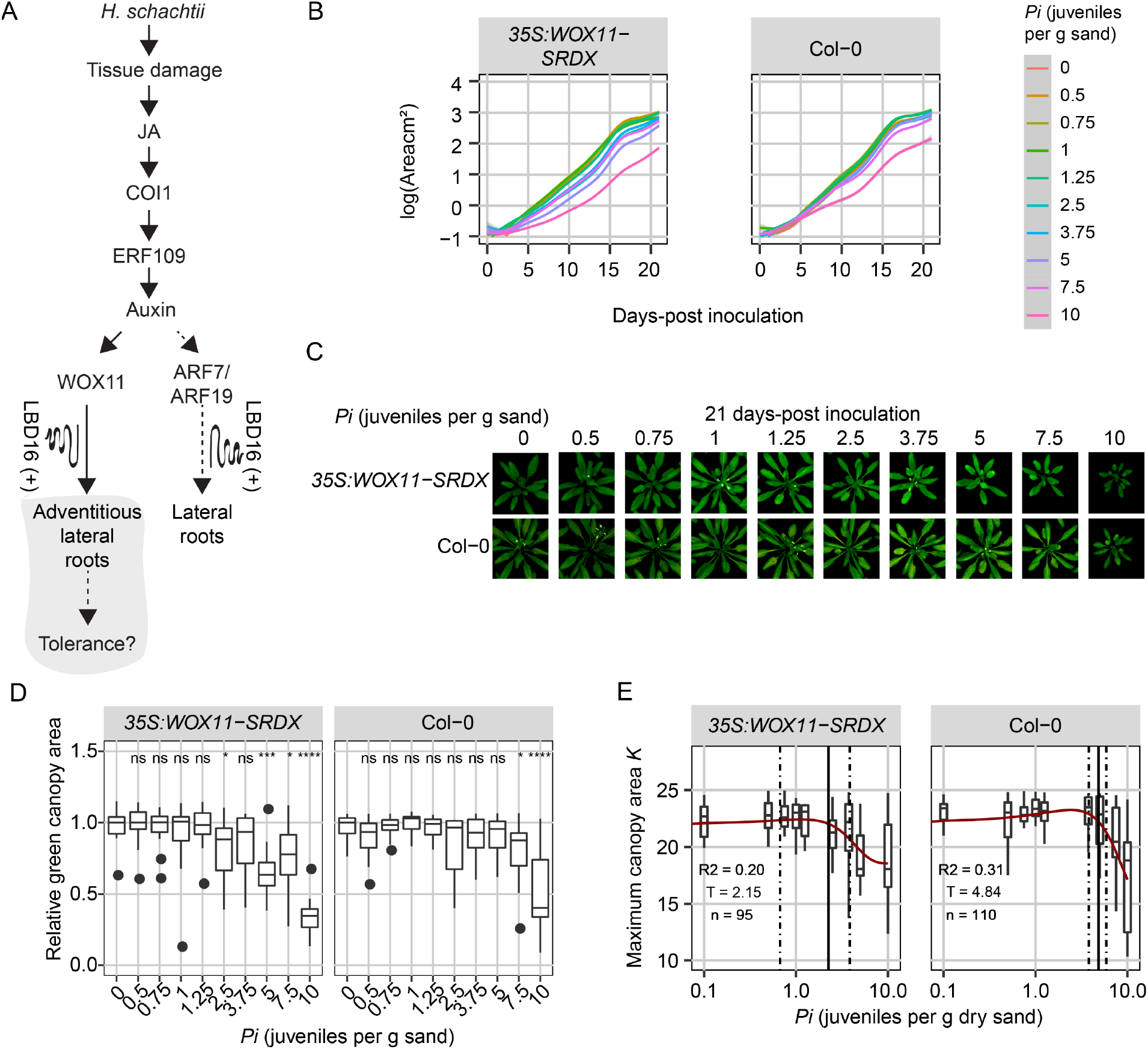
WOX11 is involved in tolerance to cyst nematode infection. **A)** Schematic diagram of *H. schachtii*-and WOX11-mediated adventitious lateral root emergence. Grey area indicates the tested part of the pathway. Curling line and ‘+’ indicate involvement of multiple proteins, including LBD16. Nine-day-old Arabidopsis seedlings were inoculated with 10 densities (*Pi*) of *H. schachtii* juveniles (0 to 10 J2s per g dry sand) in 200 mL pots containing 200 grams of dry sand. **B)** Average growth curve of Arabidopsis plants inoculated with different inoculum densities of *H. schachtii* from 0-21 dpi. Line fitting was based on a LOESS regression. **C)** Representative images of plants inoculated with *H. schachtii* at 21-days post inoculation. **D)** Relative green canopy area at 21 dpi. For the relative green canopy area, all values were normalized to the median of the measurements of the corresponding mock-inoculated plants. Data was analysed with a Wilcoxon Rank Sum test; ns= not significant, *p< 0.05, **p< 0.01, ***p<0.001 (n=10-18 plants per treatment). **E)** The maximum canopy area *K* per inoculation density of *H. schachtii*. The fitted line is from a Gaussian curve. Solid line indicates the tolerance limit. Dashed line indicates the confidence interval. *R^2^* is the goodness of the fit, *T* is the tolerance limit, and n is the number of plants used for fitting the data.

## Discussion

Excessive root branching is a classical symptom of nematode disease in plants of which the underlying causes nor the functions are well understood. Recently, we showed that endoparasitic cyst nematodes activate a JA-dependent damage signaling pathway leading to local auxin biosynthesis and subsequent *de novo* formation of secondary roots near infection sites (Guarneri et al., 2023). At the outset of this study, it was not clear if nematode-induced secondary roots emerge from primary roots following the canonical auxin-dependent pathway for the formation of acropetal lateral roots, or if they emerge following a different pathway. Our current data supports the alternative hypothesis wherein the emergence of secondary roots in response to nematode damage follows the non-canonical WOX11-dependent pathway leading to the formation of adventitious lateral roots. This induction of adventitious lateral roots near nematode infection sites compensates for the inhibition of primary root growth by root-feeding cyst nematodes. We further show that the WOX11-mediated plasticity of root system architecture contributes to the tolerance of Arabidopsis to cyst nematode infections.

Our observations demonstrate that the *de novo* root organogenesis near cyst nematode infection sites depends on WOX11, but not on ARF7/ARF19. Both WOX11- and ARF7/ARF19-mediated rooting pathways are activated by auxin, but they form a divergence point in the differentiation of adventitious lateral root primordia from lateral root primordia. WOX11 responds to auxin signals brought about by external cues, such as wounding (Sheng et al., 2017), and mediates tissue repair and regeneration mechanisms (Liu et al., 2014). In contrast, the auxin signals activating ARF7/ARF19 are thought to be developmentally regulated following endogenous rooting cues. Interestingly, both WOX11- and ARF7/ARF19-mediated root organogenesis pathways converge on LBD16 (Okushima et al., 2007; Sheng et al., 2017). Our findings indeed show that cyst nematodes induce expression of *LBD16* in a WOX11-dependent manner. However, this observation contradicts earlier work wherein *LBD16* expression was not observed in Arabidopsis infected with *H. schachtii* at similar timepoints after inoculation (Cabrera et al., 2014). It should be noted that we used a different *LBD16_pro_:LBD16-GUS* reporter line containing a much larger genomic region upstream of *LBD16* (Sheng *et al*., 2017) compared to previous studies (Okushima et al., 2007; Cabrera et al., 2014). This extended promoter region included in the *LBD16_pro_:LBD16-GUS* line harbours multiple WOX11-binding sites, which are absent in previously used *LBD16-GUS* reporter lines and which may thus explain the differences in observed *LBD16* expression in cyst nematode-infected Arabidopsis roots.

Our data further shows that both COI1 and ERF109 modulate WOX11 expression in response to cyst nematode infection, which positions WOX11 downstream of ERF109 within the JA-dependent damage signalling pathway. JA-dependent damage signaling induces local auxin biosynthesis, which drives the production of secondary roots (Guarneri et al., 2023). Auxin has been shown to directly activate WOX11 expression, and as such WOX11 connects stress-induced auxin signaling to the establishment of adventitious lateral root founder cells (Sheng et al., 2017). ERF109 most likely modulates WOX11 activity by regulating local YUCCA-mediated biosynthesis of auxin (Cai et al., 2014). However, even in the absence of ERF109 (i.e., *erf109* mutant) we observed some *WOX11-GFP* expression in nematode infection sites. This agrees with our earlier observations demonstrating that besides damage-induced local biosynthesis of auxin, auxin transported from the shoots towards nematode infection sites also contributes to local stress-induced auxin maxima (Guarneri et al., 2023). WOX11 may thus integrate local and systemic auxin-based stress response mechanisms leading to formation of adventitious lateral roots in nematode-infected Arabidopsis.

In our *in vitro* bioassays, WOX11 affected the number of secondary roots emerging from nematode-infected primary roots, but not the average secondary root length. Furthermore, we found that WOX11- mediated adventitious rooting compensated for the inhibition of primary root growth due to nematode infections, which implies that WOX11 mitigates the impact of nematode infections by adapting root system branching. This fits in the current model of wound-induced formation of secondary roots, wherein the activation of WOX11 initiates the cell fate transition of protoxylem cells into adventitious root founder cells (Liu et al., 2014). WOX11 expression is thought to be specific for adventitious root founder cells, where it activates, together with its close homolog WOX12, LBD16- and WOX5-mediated divisions to initiate the formation adventitious root primordia (Liu et al., 2014; Hu and Xu, 2016). During these divisions the expression of WOX11 decreases, because of which it affects the number of secondary roots but is less likely to alter secondary root growth.

Based on the green canopy area as a proxy for measuring the overall impact of belowground stress on plant fitness, we conclude that WOX11-mediated root system plasticity also contributes to the tolerance of Arabidopsis to cyst nematode infections. The estimated tolerance limit of *35S:WOX11-SRDX* plants for cyst nematode infections was significantly lower than for wild-type Col-0 plants. Others have shown that homologs of Arabidopsis WOX11 in rice, apple, and poplar enhance plant tolerance to abiotic stresses, such as drought and low nitrate conditions, by regulating adventitious lateral root formation (Cheng et al., 2016; Wang et al., 2020; Wang et al., 2021; Tahir et al., 2022). Furthermore, WOX11 functions as a key regulator in the regeneration of primary roots after mechanical injury by inducing the formation of adventitious lateral roots at the cut site (Sheng et al., 2017). Our study provides a first example of WOX11- mediated mitigation of the impact of belowground biotic stress.

WOX11-mediated adventitious rooting may contribute to tolerance of Arabidopsis to biotic stress by restoring the capacity of the root system to take up and transport water and minerals. Cyst nematodes modify host cells within the vascular cylinder into a permanent feeding structure, which interrupts the continuity of surrounding xylem vessels (Golinowski et al., 1996; Sobczak et al., 1997; Levin et al., 2020). As cyst nematodes develop, their feeding structures expand, consuming a larger part of the vascular cylinder while further impeding the flow of water and minerals (Bohlmann and Sobczak, 2014). This is the reason why aboveground symptoms of cyst nematodes infections are often confused for drought stress. Local and systemic auxin-based stress signals may thus activate WOX11-mediated adventitious lateral rooting to maintain the flow of water and minerals to the xylem vessels above infection sites (Levin et al., 2020). At lower inoculation densities, WOX11-meditated adventitious lateral root formation from cyst nematode infected primary roots may suffice to sustain normal Arabidopsis development and growth resulting in a more tolerant phenotype.

Recent research suggests that the cellular processes targeted by transcriptional activity of WOX11 includes the modulation of reactive oxygen species (ROS)-homeostasis. In poplar, PagWOX11/12a has been shown to regulate the expression of enzymes involved in scavenging ROS under salt stress conditions (Wang et al., 2021). In crown root meristem cells of rice, WOX11 modulates ROS-mediated post-translational modifications (i.e., protein acetylation) of proteins required for crown root development (Xu et al., 2022). ROS are required for the induction of adventitious root formation from Arabidopsis explants (Shin et al., 2022). There is also evidence that ROS modulate auxin levels during the initiation of adventitious roots from Arabidopsis explants (Huang et al., 2020). Moreover, we have recently linked tolerance of Arabidopsis to cyst nematode infections, ROS-mediated processes, and root system plasticity (Willig et al., 2022). However, further research is needed to investigate if WOX11 influences ROS-related processes, or vice versa, in infection sites of cyst nematodes in Arabidopsis roots, and if such a mechanism plays a role in WOX11-mediated root plasticity and tolerance to nematode infections.

## Supporting Information

Additional supporting information may be found in the online version of this article.

Supplemental Figure S1. Primordia formed in response to *H. schachtii* infection in *arf7-1/19-1* mutant seedlings.

Supplemental Figure S2. Root architecture comparison between *35S:WOX11-SRDX* seedlings and wild-type Col-0 seedlings.

Supplemental Figure S3. COI1 and ERF109 contribute to *WOX11* expression upon *H. schachtii* infection.

Supplemental Figure S4. Noise removal process using Gaussian Blur option in ImageJ.

Supplemental Figure S5. Adventitious lateral roots increase the total secondary root length upon nematode infection.

Supplemental Figure S6. Growth rates of *coi1-2, erf109,* and *35S:WOX11-SRDX*, and wild-type Col-0 plants over time.

Supplemental Figure S7. Growth rates of *coi1-2, erf109,* and *35S:WOX11-SRDX* plants are more affected during *H. schachtii* than wild-type.

## Funding

This work was supported by the Graduate School Experimental Plant Sciences (EPS). JJW is funded by Dutch Top Sector Horticulture & Starting Materials (TU18152). JLLT was supported by NWO domain Applied and Engineering Sciences VENI (14250) and VIDI (18389) grants. MGS was supported by NWO domain Applied and Engineering Sciences VENI grant (17282).

## Acknowledgments

We thank Prof. Viola Willemsen for providing the pWOX11::GFP, pWOX11::GFP-coi1-2, pWOX11::GFP- erf109 reporter lines, and Lin Xu for providing the 35S:WOX11-SRDX/arf7-1/19-1, arf7-1/19-1, LBD16pro:LBD16-GUS and 35S:WOX11-SRDX/LBD16pro:LBD16-GUS mutant lines.

## Author contributions

JJW, NG, JB, and GS conceived the project. JJW, NG, TvL, SW, IEAE designed and performed the experiments. MGT provided scripts for SYLM analysis. Data analysis was designed analyzed and interpreted by JJW, NG, and MGS. JJW, NG, and GS wrote the article. VW performed crosses of *coi1-2, erf109* with wildtype plants expressing *pWOX11::GFP.* VW and LX provided Arabidopsis mutant and reporter lines. VW, LX, AG, MGS, and JLLT provided critical feedback on the manuscript. All co-authors provided input for the submitted version.

## Conflict of interest

The authors declare no conflict of interest.

## Supplemental information

**Supplemental Figure S1:**
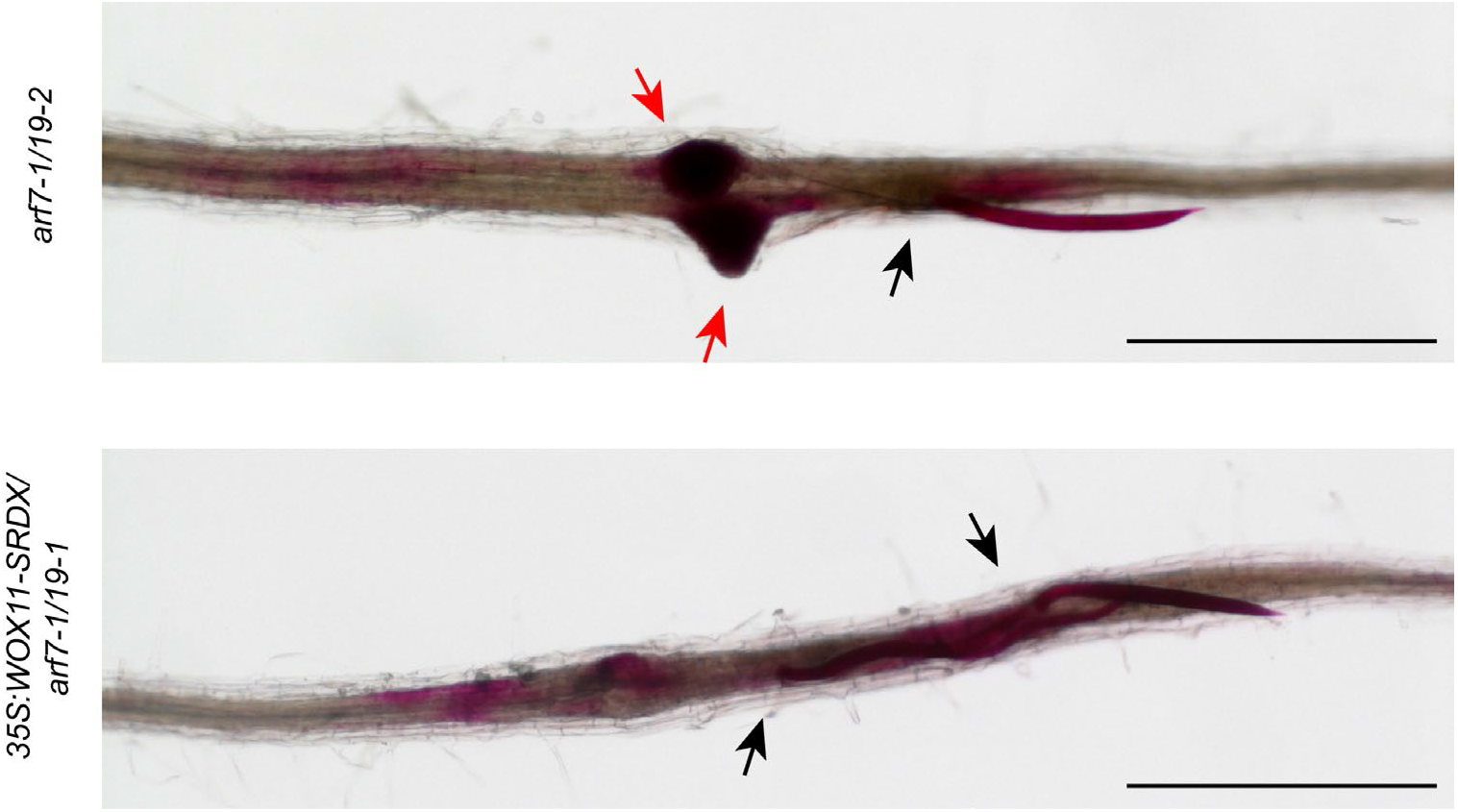
Primordia formed in response to *H. schachtii* infection in *arf7-1/19-1* mutant seedlings. Seven-day-old *35S:WOX11-SRDX/arf7-1/19-1* and *arf7-1/arf19-1* mutant seedlings were inoculated with 90 *H. schachtii* juveniles or mock inoculated. At 7 dpi nematodes were stained with fuchsin and imaged using a dissection microscope. Black arrowheads indicate head of the nematode. Red arrowheads indicate primordia. Scale bar: 500 µm

**Supplemental Figure S2:**
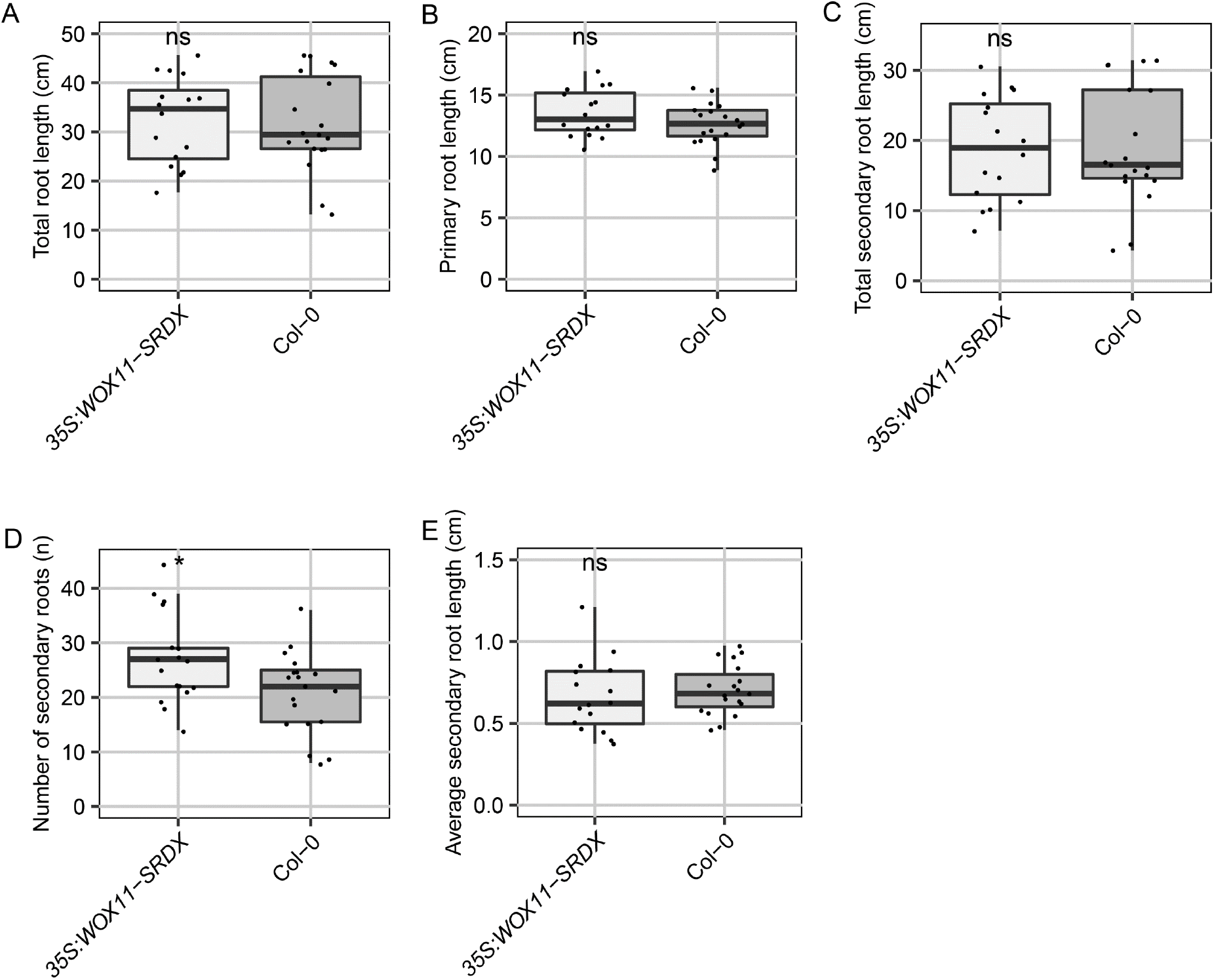
Root architecture comparison between *35S:WOX11-SRDX* seedlings and wild-type Col-0 seedlings. *35S:WOX11-SRDX* and wild-type Col-0 seedlings were grown on modified KNOP medium for 16 days. Roots were scanned and root architectural components were measured. **A)** Total root length. **B)** Primary root length. **C)** Total secondary root length. **D)** Number of secondary roots. **E)** Average secondary root length. Data from two independent biological repeats of the experiment were combined. Significance of differences between genotypes was calculated by a Unpaired Two-Samples Wilcoxon Test. (n = 14-18).

**Supplemental Figure S3:**
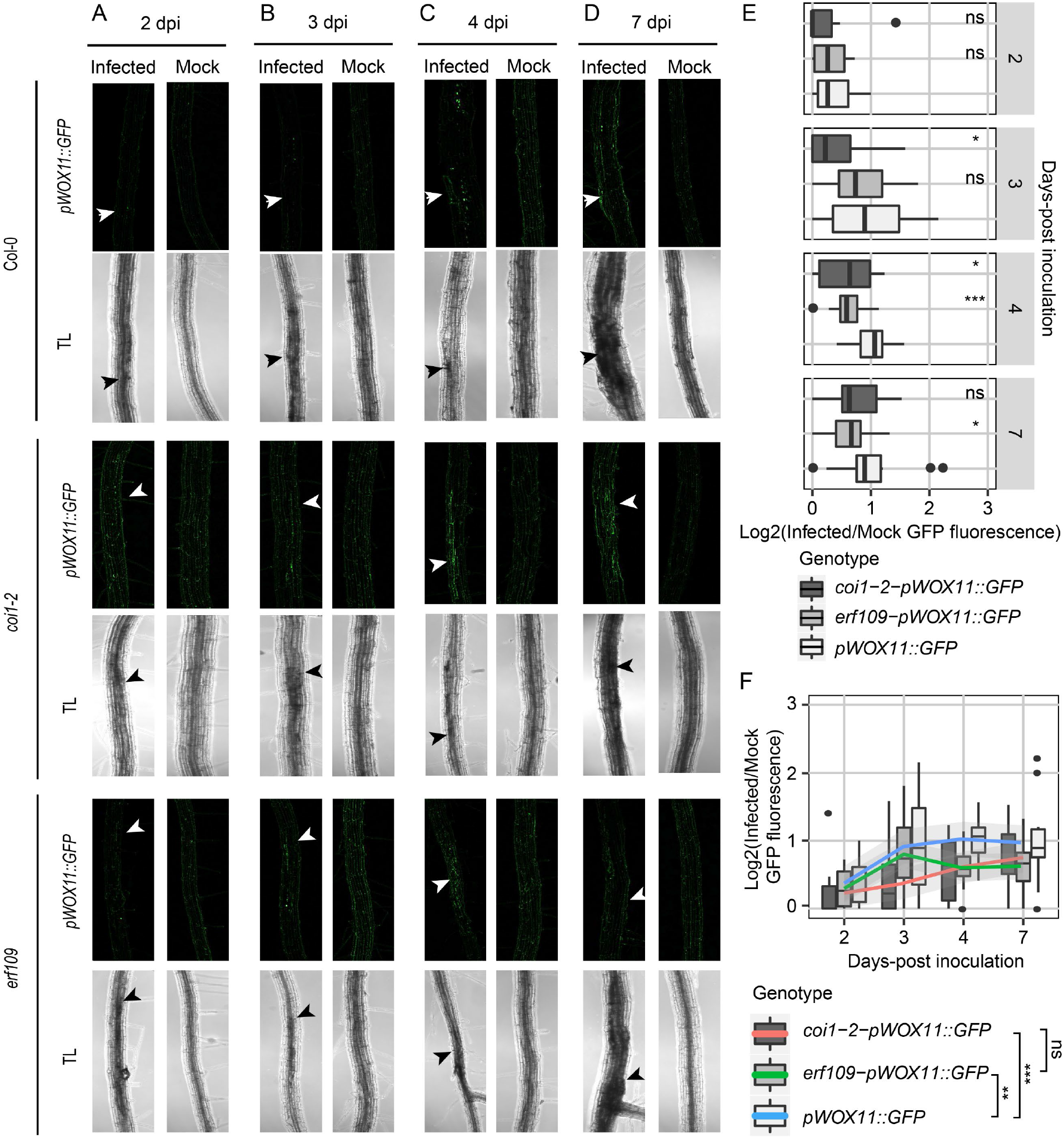
COI1 and ERF109 contribute to *WOX11* expression upon *H. schachtii* infection. A-F) Four-day-old Arabidopsis seedlings were either inoculated with 10 *H.* schachtii second-stage juveniles (J2s) or mock inoculated. At 2, 3, 4, and 7 dpi, seedlings were mounted and then imaged using a fluorescent confocal microscope. Single-nematode infection sites were selected for observation. **A-D)** Representative pictures of infected and mock-inoculated seedlings expressing the *pWOX11::GFP* construct in either wild-type Col-0, mutant *coi1-2*, or mutant *erf109* background at **(A)** 2 dpi, **(B)** 3 dpi, **(C)** 4 dpi, and **(D)** 7 dpi. To make the fluorescence more visible, the brightness was enhanced for all the representative pictures in the same way. **E)** Quantification of *pWOX11::GFP* fluorescent intensity induced by infection of wild-type Col-0, *coi1-2*, and *erf109* roots. Values represent log2 of the fluorescence ratio between the GFP integrated density of infected and noninfected roots. Data from three independent biological repeats of the experiment were combined. Significance of differences between fluorescent intensities in Co-0, *coi1-2*, and *erf109* per timepoint was calculated by a Wilcoxon Rank Sum test. ns = not significant, *p< 0.05, **p< 0.01, ***p<0.001 (n=15). **(F)** Values represent log2 of the fluorescence ratio between the GFP integrated density of infected and noninfected roots. Significance of differences between genotypes was calculated by analysis of variance. Grey area indicates the 95% confidence interval of the loess fit.

**Supplemental Figure S4:**
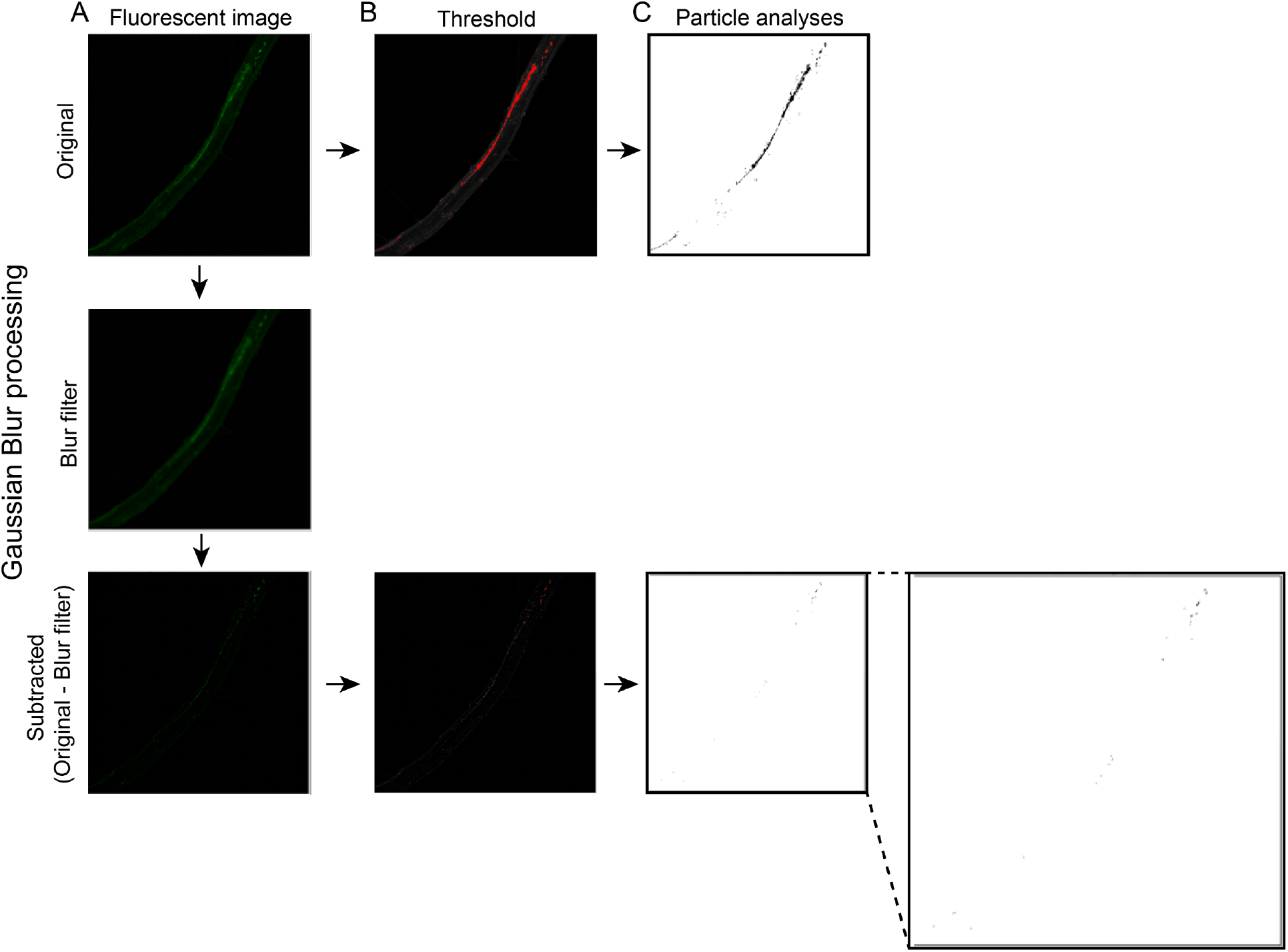
Noise removal process using Gaussian Blur option in ImageJ. **A)** The original images, which gave a lot of noise in the practical analyses (**C**) after setting the threshold (**B**) was duplicated and blurred using the Gaussian blur option in ImageJ. The blurred image was subtracted from the original image and the particles were analysed.

**Supplemental Figure S5:**
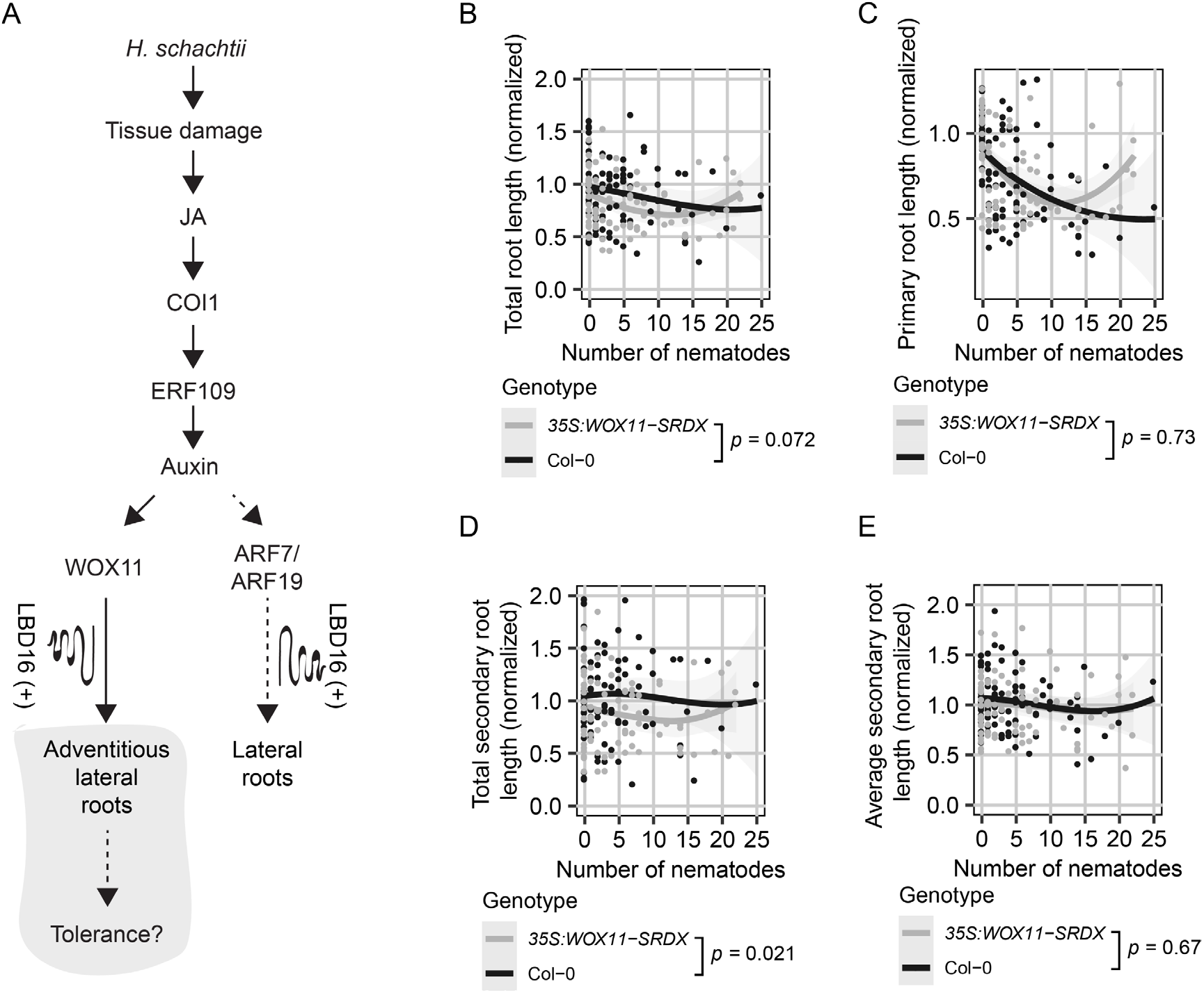
Adventitious lateral roots increase the total secondary root length upon nematode infection. **A)** Schematic diagram of *H. schachtii*- and WOX11-mediated adventitious lateral root emergence. Grey area indicates the tested part of the pathway. Curling line and ‘+’ indicate involvement of multiple proteins, including LBD16. **B-D)** Nine-day old *35S:WOX11-SRDX* and wild-type Col-0 seedlings were inoculated with densities (*Pi*) ranging from 0-7.5 *H. schachtii* J2s (per mL modified KNOP media). Roots were scanned and nematodes were counted after fuchsine staining at 7 dpi. Root architectural components of infected seedlings were normalized to the median respective component in mock-treated roots. Data of two independent biological repeats of the experiment were combined. **B)** Total root length per number of nematodes inside the roots. **C)** Primary root length per number of nematodes inside the roots. **D)** Total secondary root length per number of nematodes inside the roots. **E)** Average secondary root length per number of nematodes inside the roots. Data from two independent biological repeats of the experiment were combined. Significance of differences between genotypes was calculated by analysis of variance (n=14-18). Grey area indicates the 95% confidence interval of the loess fit.

**Supplemental Figure S6:**
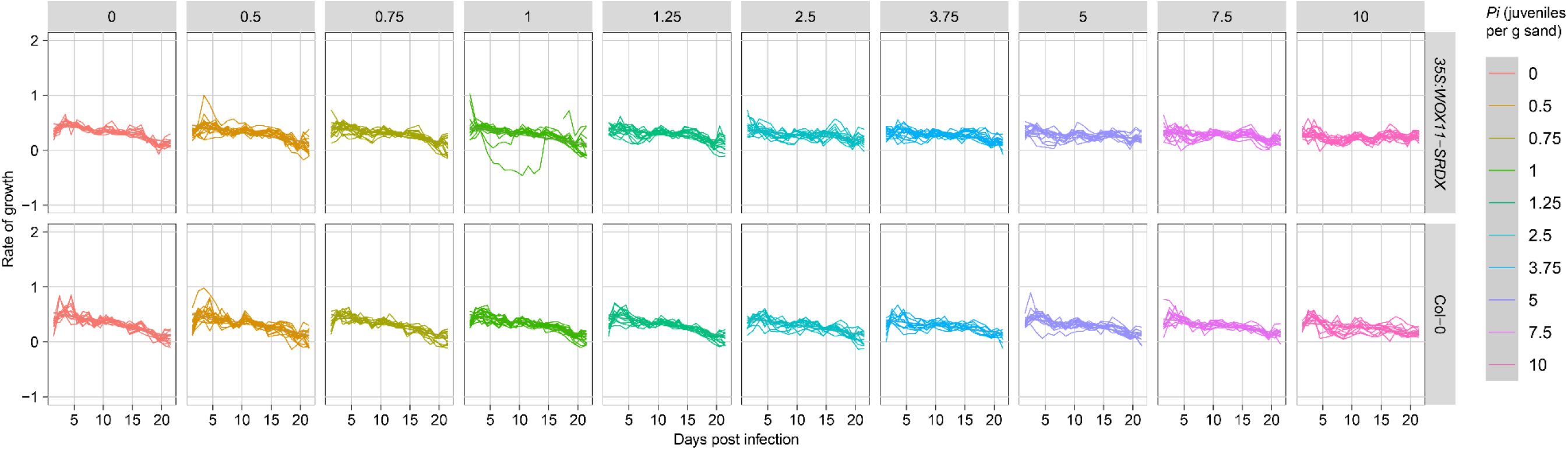
Growth rates of *coi1-2, erf109*, and *35S:WOX11-SRDX*, and wild-type Col-0 plants over time. Nine-day-old Arabidopsis seedlings (*35S:WOX11-SRDX* and wild-type Col-0) were inoculated with 10 densities (*Pi*) of *H. schachtii* juveniles (0 to 2000 juveniles per g dry sand). The growth rates of plants were calculated per day. Lines represent individual plants (n=10-18 plants per treatment).

**Supplemental Figure S7:**
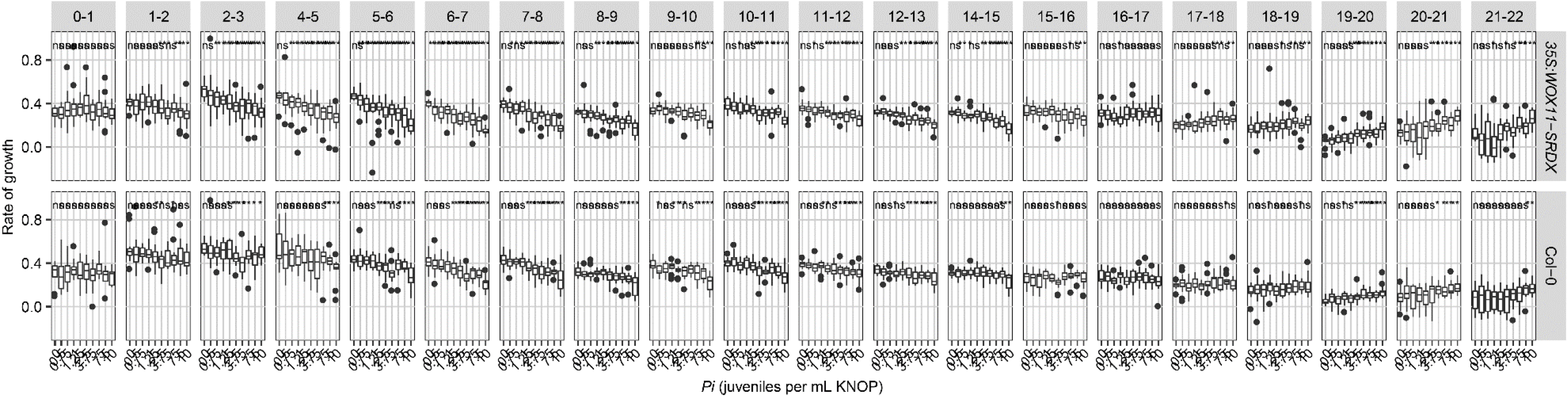
Growth rates of *coi1-2, erf109*, and *35S:WOX11-SRDX* plants are more affected during *H. schachtii* than wild-type Col-0. Nine-day-old Arabidopsis seedlings (*35S:WOX11-SRDX* and wild-type Col-0) were inoculated with 10 densities (*Pi*) of *H. schachtii* juveniles (0 to 2000 juveniles per g dry sand). The growth rates of plants were calculated per day. Boxplots represent data of x plants, the dots represent outlier measurements (1.5 times the interquartile range). Data was analysed with a Wilcoxon Rank Sum test. Ns: not significant, *p< 0.05, **p< 0.01, ***p<0.001 (n = 10-18 plants per treatment).

